# Distinct secretomes in p16- and p21- positive senescent cells across tissues

**DOI:** 10.1101/2023.12.05.569858

**Authors:** Dominik Saul, Diana Jurk, Madison L. Doolittle, Robyn Laura Kosinsky, David G Monroe, Nathan K. LeBrasseur, Paul D. Robbins, Laura J. Niedernhofer, Sundeep Khosla, João F. Passos

**Affiliations:** Division of Endocrinology, Mayo Clinic, Rochester, MN 55905, USA; Robert and Arlene Kogod Center on Aging, Mayo Clinic, Rochester, MN 55905, USA; Department of Trauma and Reconstructive Surgery, BG Clinic, University of Tübingen, Tübingen, Germany; Department of Physiology and Biomedical Engineering, Mayo Clinic, Rochester, MN 55905, USA; Robert Bosch Center for Tumor Diseases, Stuttgart, Germany; Institute on the Biology of Aging and Metabolism, Department of Biochemistry, Molecular Biology and Biophysics, University of Minnesota, Minneapolis, MN, USA

**Author notes:** **Correspondence: Dr. Sundeep Khosla Division of Endocrinology, Mayo Clinic, Mayo Clinic, 200 First Street SW, Rochester, MN 55905, USA,****, Dr. João F. Passos Department of Physiology and Biomedical Engineering, Mayo Clinic, 200 First Street SW, Rochester, MN 55905, USA**. **<u>Emails</u>:**.

## Abstract

Senescent cells drive age-related tissue dysfunction via the induction of a chronic senescence-associated secretory phenotype (SASP). The cyclin-dependent kinase inhibitors p21^Cip1^ and p16^Ink4a^ have long served as markers of cellular senescence. However, their individual roles remain incompletely elucidated. Thus, we conducted a comprehensive examination of multiple single-cell RNA sequencing (scRNA-seq) datasets spanning both murine and human tissues during aging. Our analysis revealed that p21^Cip1^ and p16^Ink4a^ transcripts demonstrate significant heterogeneity across distinct cell types and tissues, frequently exhibiting a lack of co-expression. Moreover, we identified tissue-specific variations in SASP profiles linked to p21^Cip1^ or p16^Ink4a^ expression. Our study underscores the extraordinary diversity of cellular senescence and the SASP, emphasizing that these phenomena are inherently cell- and tissue-dependent. However, a few SASP factors consistently contribute to a shared “core” SASP. These findings highlight the need for a more nuanced investigation of senescence across a wide array of biological contexts.

## INTRODUCTION

Cellular senescence is characterized by not just an irreversible cell-cycle arrest but also the development of various functional and morphological alterations in distinct cell compartments, such as the nucleus, lysosomes, mitochondria, and others^1,2^. The senescent cell-cycle arrest is largely mediated by cyclin-dependent kinase inhibitors such as p21^CIP1^ (p21 encoded by *CDKN1A)* and p16^INK4A^ (p16 encoded by *CDKN2A*)^2^. Cellular senescence also exhibits a senescence-associated secretory phenotype (SASP), which comprises a diverse array of secreted factors including immune-modulatory cytokines and chemokines, matrix remodeling enzymes, and growth factors^3^. Senescent cells play crucial roles in development, tumor suppression, and tissue repair^4–6^. However, as individuals age, the accumulation of these cells has been linked to the onset of various age-related conditions. Additionally, the removal of senescent cells either genetically or pharmacologically enhances functional outcomes in mice within the context of aging and age-related diseases, underscoring the therapeutic potential of targeting these cells^7^.

Even though there are numerous molecular changes associated with senescent cells, detecting these cells within tissues remains exceedingly challenging. Central to this difficulty is the absence of a singular specific marker for the unequivocal identification of senescent cells, as none of the markers conventionally employed in senescence detection exhibit individual specificity. Adding to this complexity, senescent cells and their SASP exhibit variability contingent upon the specific physiological context, cell type, and tissue type under investigation^8^.

The recent advancements in single-cell omics technologies offer a unique opportunity to comprehensively unravel the heterogeneity of the senescent phenotype across various cell types and tissues^9^. One of the key unresolved questions concerns the relative contributions of p16 and p21, which have been identified as critical drivers of cellular senescence, towards age-related senescence across different tissues *in vivo*. Through the analysis of multiple scRNA-seq datasets across diverse murine tissues (brain, skeletal muscle, bone, and liver) and in human skin and lung during aging, we observed that expression of p16 and p21 occurs in tissue-specific cell types which exhibit unique, often non overlapping SASP profiles, suggesting that these sub-populations have distinct functional roles. Furthermore, our findings indicate that while there are commonalities in SASP profiles in p16 and p21 expressing cells, these vary considerably according to tissue- and cell-type. However, only a small number of SASP markers can be considered as part of a “core” set. Our comprehensive analysis underscores the intricate nature of cellular senescence and the SASP, emphasizing the important role of single-cell studies to fully elucidating and characterizing senescence in aging tissues.

## RESULTS

### Unraveling p16- vs. p21-associated SASP in the murine brain

Previous studies have shown that markers of cellular senescence increase during aging in murine brain and, importantly, that clearance of p16+ cells enhances cognitive function in aged mice^10^. To further investigate cellular senescence in the brain, we conducted an in-depth analysis of scRNA-sequencing datasets^10^ to profile and compare the cellular composition and transcriptomes of young (4m) and old mouse (24m) hippocampi; a brain region known for its involvement in memory formation. Our analysis initially identified five primary cell populations within the hippocampus (**Fig. 1a**). To mitigate potential confounding factors introduced by inflammatory immune cells, we refined our focus by excluding CD45^high^ cells. This criterion still allowed the inclusion of microglia in our analysis which are characterized by CD45^low/intermediate^ expression^11^. Further filtering steps involved ensuring the absence of the proliferation marker Ki67 (*mKi67*) and verifying that the selected cells were not in the S phase since senescent cells are arrested in G1/2 phases of the cell-cycle^12^. Subsequently, p16(*Cdkn2a*)-positive cells and p21(*Cdkn1a*)-positive cells were isolated. Among the p16+ cells, microglia and oligodendrocytes appeared as the main subpopulations, whereas in the p21+ cells, microglia predominated (**Fig. 1b**).

This analysis revealed the presence of 3 subpopulations, 1) cells exclusively expressing p21; 2) cells exclusively expressing p16; and 3) cells expressing both p21 and p16 (**Fig. 1c, d**). To gain deeper insights into the composition of the SASP within these identified subpopulations, we leveraged our previously established SenMayo gene set^13^, which has been demonstrated to be commonly regulated across various age-related transcriptome datasets. We observed a distinct SASP profile for p21+ and p16+, characterized by a limited overlap in SASP gene expression. Notably, only two genes, *Cxcl16* and *Plaur*, were found to be expressed in both subpopulations (**Fig. 1e**). Likewise, when we analyzed an extensive list of secreted proteins not limited to the known SASP^14^ plus SenMayo, we observed a limited overlap in gene expression between p21+ and p16+ cells. Only four genes, *Col8a2*, *Plaur*, *Cxcl16* and *Fbln5* were found to be shared between these two subpopulations (**Extended data Fig. 1**).

**Figure 1:**
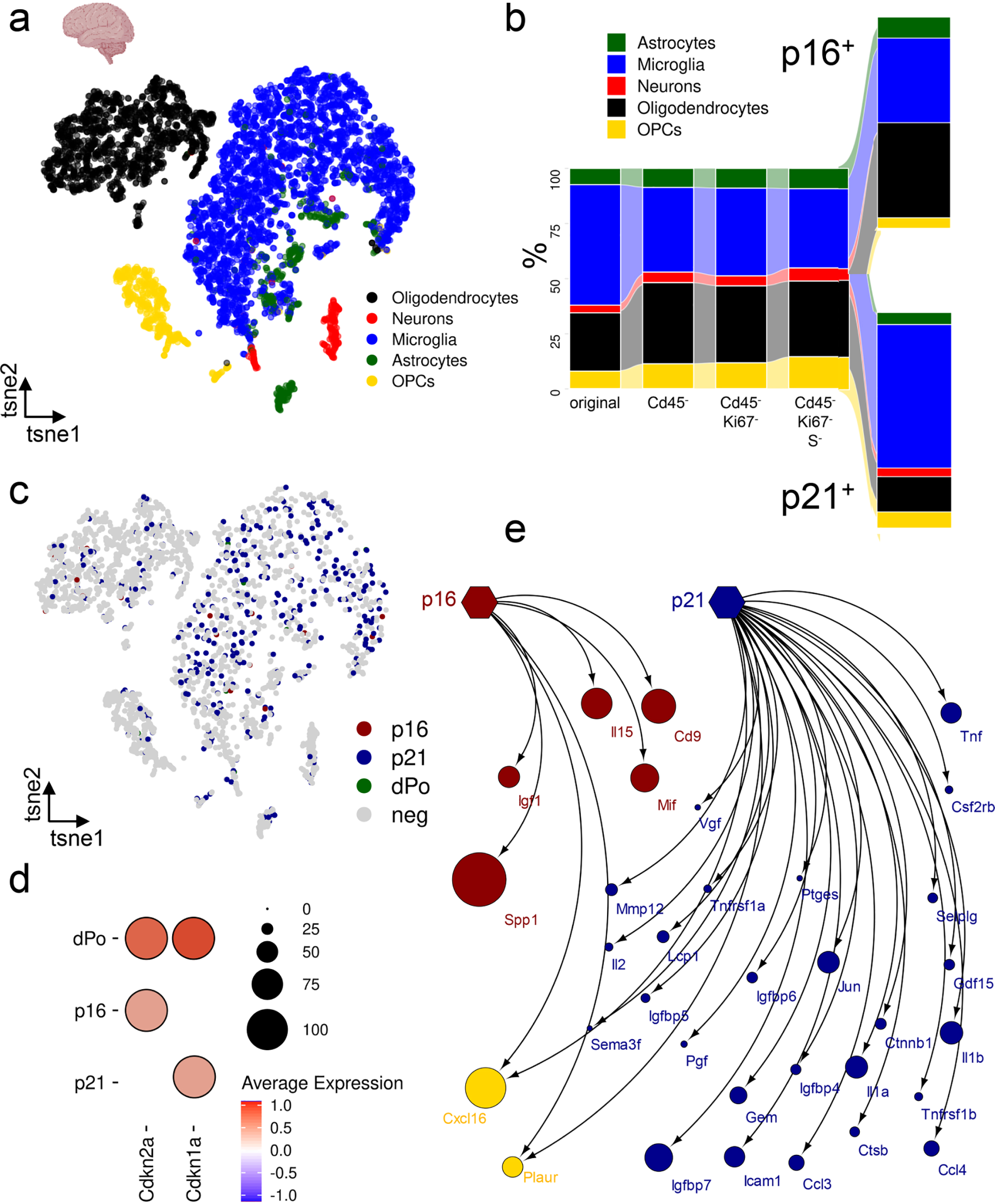
Distinct p16+ and p21+ cell populations with differential SASPs in the murine hippocampus. (**a**) A tSNE plot depicting the five main populations of young and old murine hippocampus (PMID: 33470505, GSE161340). (**b**) From the original population, CD45-cells are selected, followed by Ki67-negativity and cells not in the S phase. From these cells, just p16(*Cdkn2a*)+ cells and p21(*Cdkn1a*)+ cells are selected. From the p16+ cells, microglia and oligodendrocytes depict the main populations, while in the p21+ population, microglia is predominant. (**c**) tSNE visualization of p16 (red) and p21 (blue) cells, along with few double-positive cells (green), shown in the Cd45-Ki67-S-population. (**d**) The dPo (double positive) cells are high in both Cdkn2a and Cdkn1a while p16+ cells just express Cdkn2a, but no Cdkn1a and vice versa. (**e**) Utilizing the SenMayo gene set, SASP factors exclusively expressed by p16+ cells are fewer than those expressed in p21+ cells, with some (Cxcl16, Plaur) being expressed by both. Very few SASP genes are expressed by p16-negative and p21-negative cells. The size of the dots represents the fold change compared to all other populations shown in (c).

Overall, these data emphasize the diversity as well as variability in SASP profiles among cells with core features of senescence in the brain.

### Comparing p16 and p21-associated SASP across murine tissues

After our initial observations in the brain, we extended our investigation to assess the generality of our findings in diverse murine tissues during aging. We specifically focused on skeletal muscle, bone, and liver, as previous research had indicated age-dependent rises in senescence-associated markers and demonstrated the benefits of eliminating senescent cells for the functional outcomes of these organs^15–17^. We used single-cell RNA-seq datasets comparing young and old mice. In total, we successfully identified 9, 14, and 13 distinct cell populations in skeletal muscle (**Fig. 2a**), bone (**Fig. 2d**), and liver (**Fig. 2g**), respectively. We then followed a similar methodology as in our brain analysis, excluding cells that were positive for Ki67 and in the S-phase of the cell cycle, as well as CD45^high^ cells.

In our examination of all three tissues (**Fig. 2**), we observed a consistent pattern. We identified distinct cell subsets that exclusively expressed either p21 or p16, and a rare subgroup of cells that co-expressed both markers (**Fig. 2b, e & h**). Notably, in all three tissues, the population of cells expressing p21 was considerably more abundant. Our analysis utilizing the SenMayo dataset revealed that the trend found in brain extended across tissues: a limited set of SenMayo factors were uniquely expressed by p16+ cells, while p21+ cells exhibited a broader range of SenMayo factors (**Fig. 2c,f & i**). Although a small number of genes were co-expressed by cells positive for both p16 and p21, these were not consistently shared among the three tissues. Indeed, our investigations suggest that there is a distinctiveness in the composition of SASP profiles for each tissue, irrespective of the expression of p21 and p16. If we extend our analysis to a larger set of genes including an extensive list of secreted proteins plus SenMayo, we observe a similar pattern across murine tissues with p21+ cells being associated with a more abundant secretory profile and a limited overlap between the secretory profile of p21+ and p16+ cells (**Extended Data Fig. 2a-c**).

**Figure 2:**
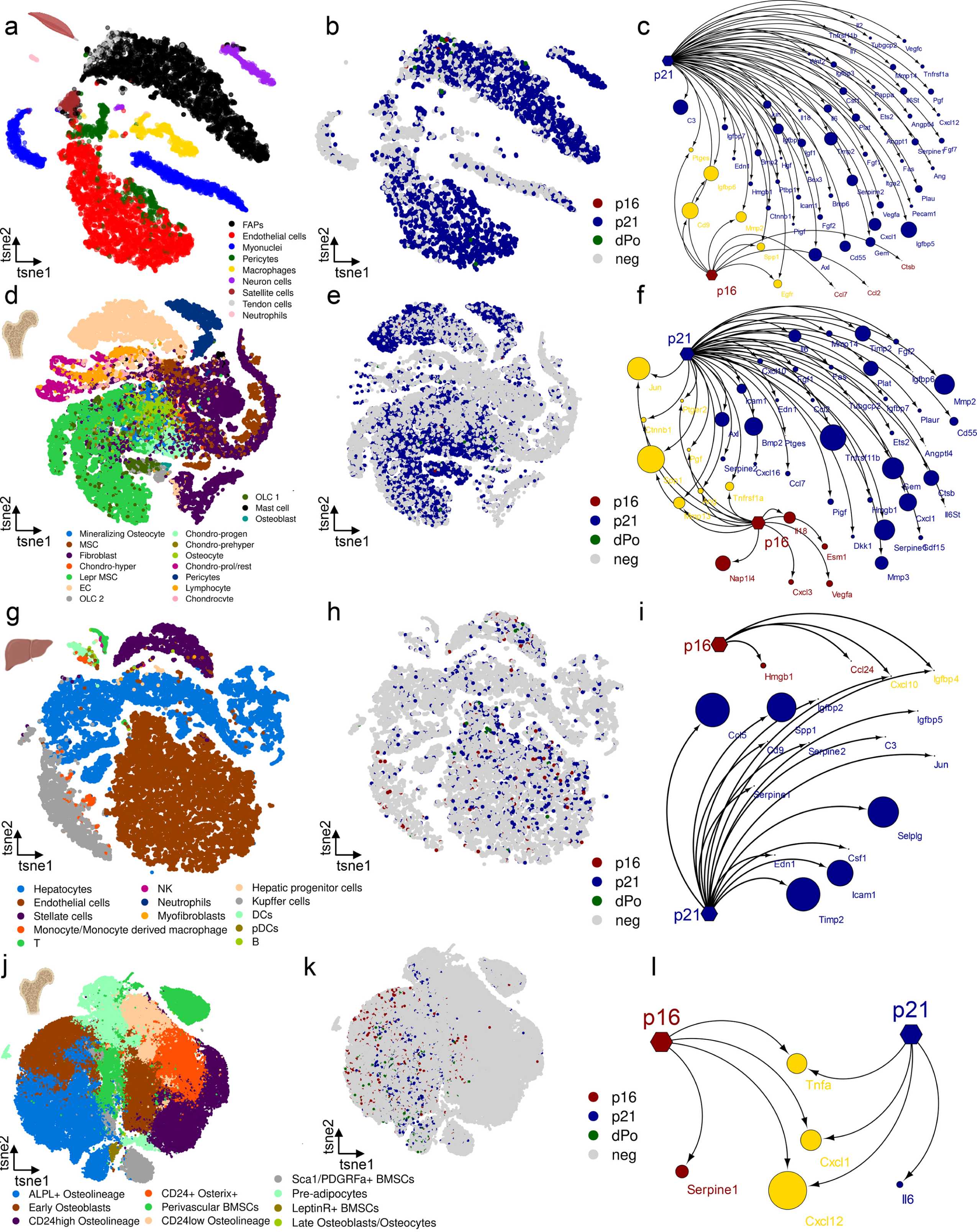
Comparative analysis of p16+ and p21+ cells and their SASP in different murine tissues during aging. (**a**) In murine skeletal muscle (PMID: 36147777, GSE172410), nine different cell types can be distinguished. (**b**) p21+ cells constitute the majority, while double positive (dPo) cells are infrequent. (**c**) The SASP is heterogeneous, with p21+ cells expressing a vast array of SASP factors, exhibiting minimal overlap with p16-associated SASP factors. (**d**) In murine bone (PMID: 31130381, GSE128423), 17 cell types can be distinguished. (**e**) p21+ cells once again dominate the senescent cell population, and the (**f**) SASP in murine bone remains diverse, with p21+ cells expressing a larger number of SASP factors compared to p16+ cells. (**g**) In the murine liver (PMID: 34755088, GSE166504), 13 different cell types are identified. (**h**) The liver has the lowest proportion of p21+ cells from all tissues analyzed, although they still form the majority of senescent cells, with a few dPo cells. (**i**) The p21-associated SASP is larger compared to the p16-associated SASP, with limited overlapping SASP factors. (**j**) In a single-cell proteome dataset analyzed by CyTOF in bone (PMID: 37524694), 10 different cell types are distinguished. (**k**), p21+ cells are present in a higher proportion than p16+ cells, and double-positive cells are rare. (**l**) The secretome exhibits unique factors for p16+ and p21+ cells, with overlap in Cxcl12, Cxcl1, and Tnf.

**Figure 3:**
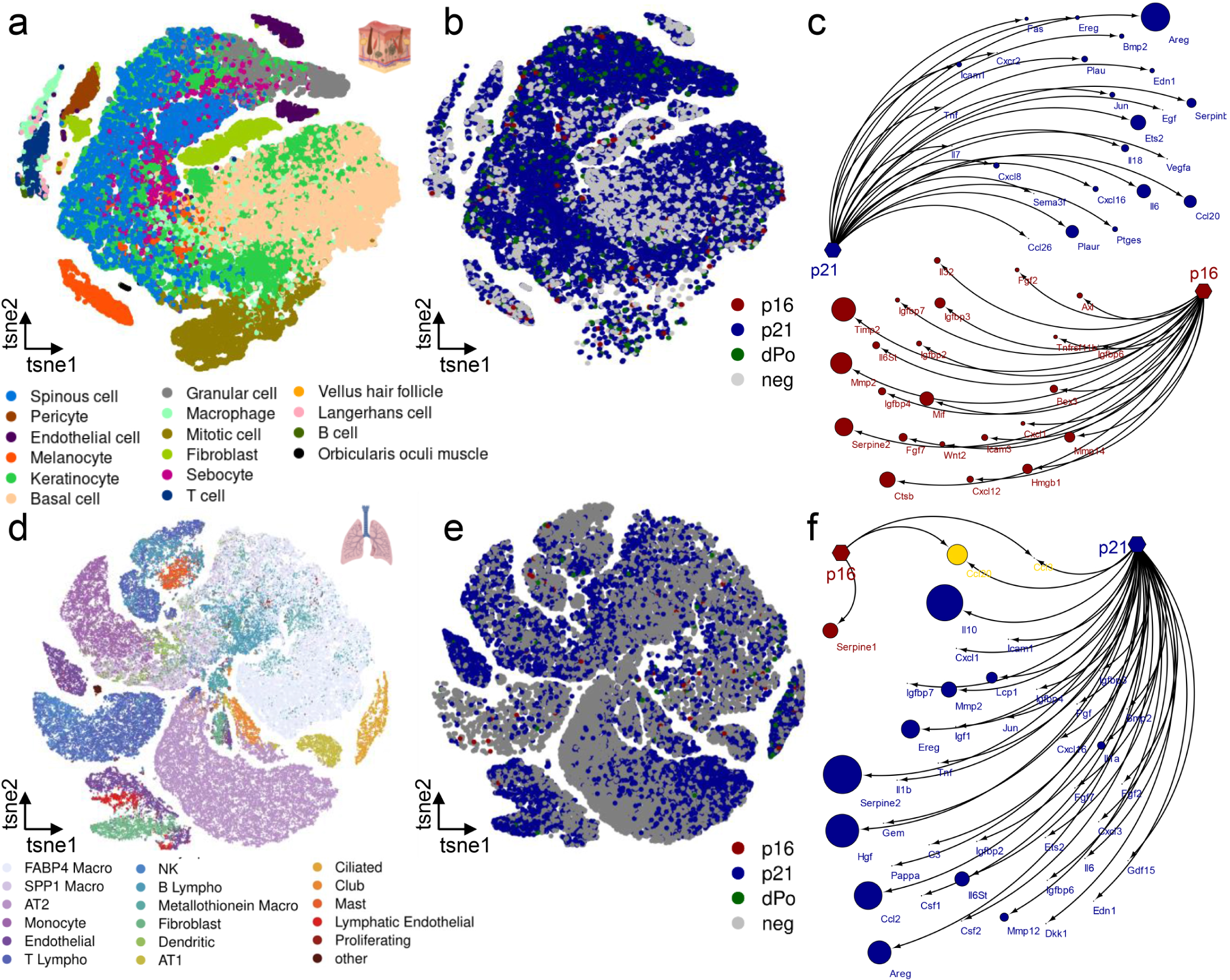
Comparative analysis of p16+ and p21+ cells and their SASP in human skin and lung during aging. (**a**) In human skin samples (PMID: 33238152, HRA000395), 16 distinguishable cell populations are identified. (**b**) Among these populations, p21+ cells are notably abundant, while double-positive (dPo) cells are rare. (**c**) The SASP expressed by p21+ cells is significantly larger compared to the SASP associated with p16+ cells. (**d**) In the human lung (PMID: 37706427, GSE122960, GSE128033, GSE130148, GSE212109), 18 discernible cell types are identified, and (**e**) within these cell types, p21+ cells constitute the majority of senescent cells. (**f**) The SASP profile of p21+ cells is extensive compared to that of p16+ cells, with minimal overlap.

This observation aligns with transcriptomics data from a previous study, where the overexpression of p21 and p16 in mouse embryonic fibroblasts (MEFs) resulted in distinct SASP profiles^18^. Certain factors were unique to either p21 or p16 overexpressing senescent cells, with some overlap (**Extended Data Fig. 3**).

Furthermore, to investigate if our findings extend to the protein level, we conducted an analysis using data derived from mass cytometry by time-of-flight (CYTOF) conducted on the bones of young and aged mice^19^. This technique provided a detailed examination of cell-type and senescence-associated protein markers at the single-cell level (**Fig.2 j**). Similar to the scRNAseq analysis, we excluded cells positive for Ki67 and CD45. Our analysis revealed distinct subsets of cells expressing either p21 or p16 exclusively, along with a rare subgroup co-expressing both markers at the protein level (**Fig.2 k**). It is important to note that CYTOF methodology relies on targeted panels of antibodies, which limited our analysis of SASP components. Nevertheless, we identified proteins such as Serpine1 exclusively expressed in p16+ cells and Il6 exclusively expressed in p21-positive cells, reinforcing the existence of distinct SASP profiles (**Fig. 2 j-l**).

### Comparing p16- and p21-dependent SASP in human tissues during aging

Subsequently, we aimed to assess the consistency of our findings across species. To achieve this, we analyzed scRNA-seq datasets from human skin and lung tissues during aging^20,21^. To examine human skin, we utilized a recently published dataset in which scRNA-sequencing was conducted on human eyelid skin samples from individuals spanning an age range of 18 to 76 years^20^. Here, we identified 16 distinct cell populations (**Fig. 3a**). We then proceeded to remove Ki67+, CD45^high^ and S-phase cells and observed three subpopulations: p21-exclusive, p16-exclusive, and p21-p16 co-expressing cells, with p21-exclusive being the predominant group (**Fig. 3b**). Interestingly, by utilizing the SenMayo dataset, we identified no overlap between p21 and p16 expressing cells, underscoring their distinctiveness as separate cell populations (**Fig. 3c**). A similar pattern was observed in human lung. Here we utilized a recently published scRNA-seq dataset from healthy human lungs ranging from 21 to 78 years of age^21^. Within our analysis, we identified 18 distinct cell populations (**Fig. 3d**). After excluding Ki67+, CD45^high^ and S-phase cells, we observed that a significant majority of cells displayed p21 expression, while only a small number of cells exhibited p16 or the combination of p16 and p21 (**Fig. 3e**). Notably, cells that were p16+ demonstrated limited expression of SenMayo components, with *Serpine1* being the exclusive gene expressed by this subgroup. In contrast, p21+ cells exhibited the expression of multiple SenMayo components (**Fig. 3f**). When we broaden our examination to include a wider range of genes which involves a comprehensive list of secreted proteins^22^ alongside SenMayo^13^, we observed that there are relatively few SASP/secreted protein genes that are common between p21 and p16 expressing cells in both human skin and lung (**Extended Data Fig. 4a & b**). This further underscores the functional specificity inherent in these sub-populations.

### Common p16 and p21 SASP across murine and human tissues

Following our identification of significant heterogeneity in the SASP genes expressed between p16+ and p21+ senescent cells among tissues in both murine and human subjects, we sought to investigate which elements exhibit commonality across diverse tissues and species (**Fig. 4a and b).** When investigating the p16+ SenMayo gene set, we noticed that only a portion of SenMayo genes were expressed in two or more of the analyzed tissues. This subset included *Spp1*, *Cd9*, *Mif*, *Ctsb*, *Mmp2*, *Igfbp6*, *Hmgb1* and *Igfbp4* (**Fig. 4c**).

In the case of p21, we observed that *Icam1* and *Jun* were consistently expressed across all six analyzed tissues, while several other factors, including *Cxcl16*, *Il6*, *Pgf*, *Ets2*, *Igfbp4*, *Igfbp6*, *Igfbp7*, *Bmp2*, *Gem*, *Serpine2*, *Ptges*, and *Edn1*, were expressed in four or more of the six analyzed tissues (**Fig. 4d**). Combining p16 (red) and p21 (blue) markers that are expressed in more than two (p16) and four (p21) tissues, a “core” SASP comprised of *Jun, Igfbp4, Igfbp6 and Spp1* is established. While *Jun* may exhibit a stronger association with p21, *Igfbp4* and *Igfbp6* are prominently expressed by both cell types, suggesting these as “common” SASP factors (**Fig. 4e**). In our search for a common SASP signature, we broadened our analysis to encompass a greater number of secreted factors, in conjunction with SenMayo. This approach enabled us to identify 16 genes expressed in p16+ cells that were consistently present in (at least) 3 of the analyzed tissues (**Fig. 4f**). In p21+ cells, we identified 50 distinct genes expressed in a minimum of four tissues, among which 5 genes were expressed in five tissues, and a single gene, *Jun*, was expressed in all analyzed tissues (**Fig. 4g**). Interestingly, this factor, along with *Edn1*, has been previously identified by SenMayo. In contrast, *Col18a1*, *F3*, *Nampt* and *Sdc4* are expressed across five tissues and are not associated with SenMayo. Upon combining all factors to identify a “common” SASP, we observed that *Jun* remains abundant primarily in p21+ cells. Additionally, *Sparc* is expressed in 3 out of 6 tissues by p16+ cells and in 4 out of 6 tissues by p21+ cells, thereby establishing it as another “common” SASP marker. (**Fig. 4h**).

**Figure 4:**
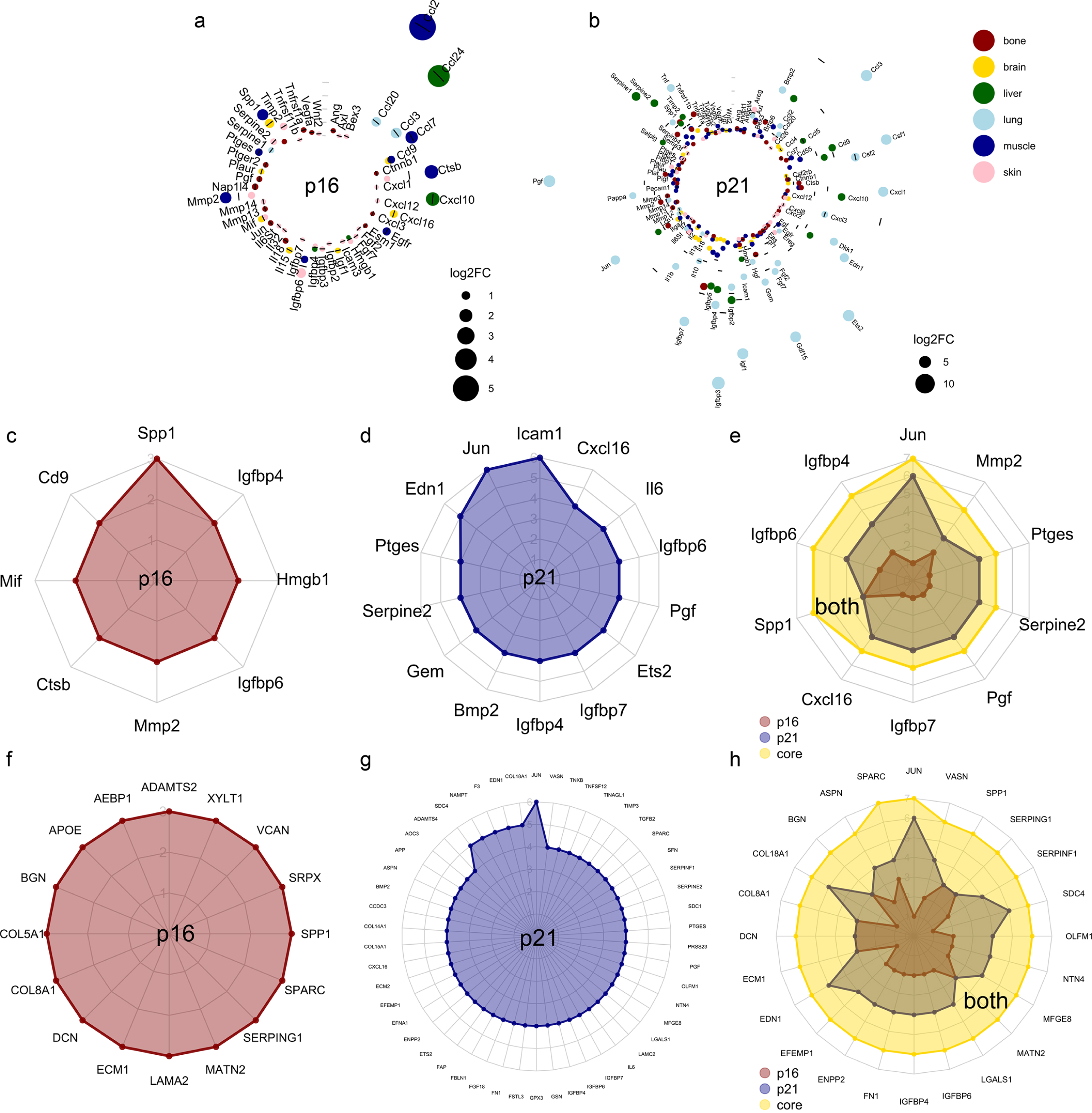
Common p16- and p21-associated SASP across murine and human tissues. (**a**) The p16-associated SASP is characterized by the predominant presence of *Ccl2* and *Ccl24*, with some markers, such as *Spp1*, expressed in multiple tissues. (**b**) The p21-associated SASP displays greater diversity, with genes like *Icam1* and *Jun* commonly expressed across tissues. (**c**) From the SenMayo panel, the p16-core-SASP includes *Spp1*, *Cd9*, *Mif*, *Ctsb*, *Mmp2*, *Igfbp6*, *Hmgb1* and *Igfbp4*. (**d**) The p21-associated SASP is dominated by *Icam1* and *Jun* that are expressed consistently by p21-positive cells in all six analyzed tissues. (**e**) A “core”-SASP for senescent cells comprises *Jun*, *Igfbp4*, *Igfbp6* and *Spp1*. (**f-h**) The secretome associated with both p16 and p21, derived from the SenMayo and whole secretome, exhibits a higher number of shared markers across various tissues. *The depicted genes are those significantly overexpressed (p<0.05, FC>0), with cell type colors corresponding to those in Fig. 1–3. In (a) and (b), the dot size reflects the log2FC. For (c), only markers expressed in 2 or more tissues are shown; in (d), those expressed in 4 or more tissues; in (f), those expressed in 3 tissues; and in (g), those expressed in more than 4 tissues are displayed*.

### Heterogeneous intercellular communication highlights the functional diversity of p21+ and p16+ cells

The SASP plays a crucial role in intricate intercellular communication, exerting complex effects on neighboring cells. These effects include the propagation of senescence^23^, modulation of tissue repair processes^6,24,25^, and recruitment of immune cells^26^. We sought to determine if senescence sub-types (p21+ and p16+ cells) not only differed in their transcriptomes, but also in how they communicate with other cells. Thus, we utilized CellChat, a commonly used tool for inference of cell-cell communication^27^. Our analysis focused on both p21+ and p16+ cells and their interactions with neighboring cell types across the mentioned tissues. An initial pairwise examination of the communication patterns of these specific p21+ or p16+ cells within each tissue shows a heterogeneity for the most important 17 signaling pathways (**Fig.5a**). A more detailed examination in each tissue shows that in brain (**Fig. 5b**), p21+ cells mostly used the CCL and JAM-pathway for communication with microglia, while p16+ cells favored the MAG-pathway to communicate with oligodendrocytes. In bone (**Fig. 5c**), p21+ cells used the THBS- and FN1-pathway exclusively to communicate with hypertrophic chondrocytes and osteolineage cells, while p16-positive cells utilized other mechanisms. In skin (**Fig. 5d**), the CD99-pathway was favored by p16+ cells and the DESMOSOME-pathway by p21+ cells to communicate to granular and spinous cells as sebocytes. Muscle (**Fig. 5e**) was characterized by a low number of p16+ cells and a high secretory pattern of the p21+ cells, which mostly used the LAMININ pathway to communicate with Fibro-Adipogenic Progenitors (FAPs) and tendon cells. In liver (**Fig. 5f**), p21+ cells favored the JAM pathway to communicate with myofibroblasts and hepatic progenitors, while p16+ cells used the CDH5-pathway for communicate with endothelial and Kupffer cells. In lung (**Fig. 5g**), the FN1-pathway was more used by the p16+ cells, while p21+ cells favored the MIF-pathway. These findings emphasize that the diversity of the p21 and p16-dependent SASP across different tissues reflects the intricate nature of intercellular communication.

**Figure 5:**
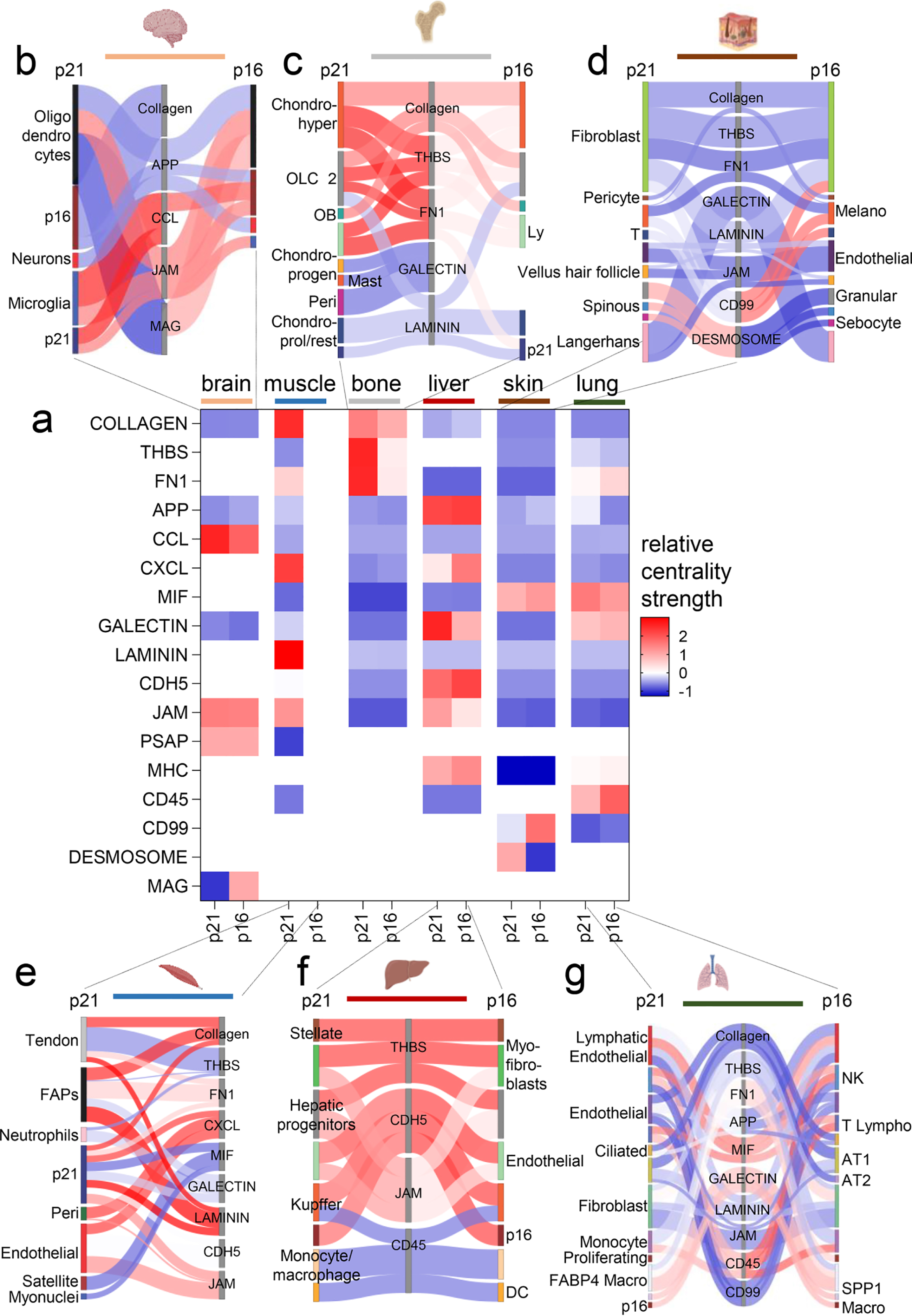
Unequal communicational patterns between p21+ and p16+ cells across tissues. (**a**) The network centrality scores depict low (blue) and high (red) communication networks for seventeen significant interactions, with p21+ and p16+ cells shown pairwise in each tissue. (**b**) In the brain, p21+ cells exhibit higher interaction strength in CCL communication, primarily to microglia cells, while p16+ cells demonstrate increased secretory activity via MAG to oligodendrocytes. (**c**) In bone, p21+ cells exclusively utilize the THBS and FN1 pathways to communicate with hypertrophic chondrocytes, osteoblasts, and lymphocytes. (**d**) In the skin, p16+ cells show higher interaction strength in the CD99 pathway to fibroblasts and melanocytes, while p21+ cells predominantly signal through the DESMOSOME pathway to granular and spinous cells, including sebocytes. (**e**) In muscle, p21+ cells mainly signal through the COLLAGEN and LAMININ pathways to fibro-adipogenic progenitors (FAPs) and tendon cells, as p16-positive cells are too few for calculating communicational patterns. (**f**) In the liver, the JAM pathway is predominantly used by p21+ cells to communicate with myofibroblasts and hepatic progenitors, whereas p16+ cells primarily employ the CDH5 pathway to interact with endothelial and Kupffer cells. (**g**) In the lung, p16+ cells exhibit the highest signaling strength in the CD45 pathway, mainly used to communicate with NK cells and T lymphocytes, while p21+ cells primarily use the MIF pathway for contact with lymphatic endothelial and NK cells. *Displayed are only significant interactions (padj<0.05), with cell type colors corresponding to those in* Figures 1–3*. The color scale (blue-white-red) of the interactions in (B-G) corresponds to the centrality score in (A)*.

This heterogeneity becomes apparent when exploring the transcription factors that control gene expression in p16+ and p21+ cells. To achieve this, we employed SCENIC, a computational method that allows the prediction of interactions between transcription factors and target genes based on single-cell RNA-seq data^28^. SCENIC analyses reveal that transcription factors regulating gene expression in p16 (**Extended data Fig. 5a & b**) and p21 (**Extended data Fig. 5c & d**) positive cells are mostly tissue specific, with minimal overlap between tissues. This implies that the SASP is not exclusively specific to p16+ or p21+ cells; rather, these senescent subtypes are further transcriptionally regulated by factors specific to the tissue in which they are situated. In summary, the diversity in cellular communication and transcriptional regulation between p16+ and p21+ cells within the unique environments of the six analyzed tissues underscores their heterogeneity and suggests potentially distinct functions.

## Discussion

p21 and p16 are both associated with the induction of senescence, but they do not always occur together in senescence. Multiple studies *in vitro* have shown that their presence and expression can vary depending on the senescence-inducing stimuli, the cell type, and the specific context^29^. p21 is a direct target of the tumor suppressor protein p53. When DNA damage or other stressors occur, p53 becomes activated and binds to the p21 promoter, leading to increased p21 expression. p21, in turn, inhibits the activity of cyclin-dependent kinases (CDKs), halting the cell cycle and promoting senescence. p16, on the other hand, acts through the Retinoblastoma (Rb) pathway^30^. p16 inhibits CDK4 and CDK6, which are responsible for phosphorylating Rb. When Rb is not phosphorylated, it remains active and prevents the cell from progressing through the cell cycle. Both pathways are intricate as they involve numerous upstream regulators and downstream effectors, as well as the presence of diverse side branches. Furthermore, these pathways exhibit substantial interconnections and crosstalk^31^.

p16 and p21 are widely employed as the two most common markers for identifying senescent cells. Their bulk expression has been extensively utilized to identify senescent cells in different tissues affected by aging or other pathological conditions^2^. However, only recently, with the advancement of single-cell omics technologies, can we truly examine the full extent of their heterogeneity *in vivo*^32^.

Our study clearly demonstrates that, in various aging tissues, cells expressing p21 and p16 at both the transcript and protein levels frequently constitute separate subpopulations characterized by distinct SASP compositions. This suggests the activation of these two pathways may result in functionally diverse consequences and is reflected in the engagement of different intercellular communication pathways. Nevertheless, it is plausible that some of these cells are at varying stages of senescence induction, with p21 being triggered earlier following exposure to a stressor and p16 being induced in a later stage of senescence as previously shown *in vitro*^33^. Consistent with the functional diversity of these sub-populations, a recent study has demonstrated that p21 and p16 overexpression elicit distinct SASPs, with p21’s SASP promoting immunosurveillance^18^. p21 upregulation has also been suggested as a mechanism that enables senescent cells to resist apoptosis thereby facilitating their retention in tissues^34^. Moreover, recently developed transgenic models where either p21+ or p16+ cells can be cleared have been shown to have different functional outcomes. For example, selective elimination of p21+ senescent cells, as opposed to p16+ ones, effectively prevents radiation-induced osteoporosis^35^.

Interestingly, the composition of SASP in cells expressing p21 and p16 exhibited significant variation across different tissues, with only a limited number of common SASP factors. This observation holds significant conceptual implications. It indicates that the phenotypes resulting from the activation of senescence-associated pathways during aging are strongly influenced by the specific cell type involved. Moreover, it raises the intriguing possibility that distinct intrinsic mechanisms may contribute to senescence in various tissue types during aging. Finally, it suggests that analysis of SASP components in tissues should be comprehensive in nature and conducted at single-cell resolution.

Our findings hold significant implications for the implementation of senolytic therapies in clinical contexts. They underscore the importance of adopting a context-specific approach not just taking into consideration the subtype of senescent cells, but also the tissue which is being targeted.

## Materials and methods

### Single-cell analysis

The scRNA-seq data were aligned and quantified using the 10X Genomics Cell Ranger Software Suite (v6.1.1) against the murine reference genome (mm10) and human reference genome (hg19). The Seurat package (v4.3.0.1 and 5.0.0) (PMID: 29608179, PMID: 31178118) was used to perform integrated analyses of single cells. Genes expressed in <3 cells and cells that expressed <200 genes and >20% mitochondria genes were excluded from downstream analysis in each sample. The datasets were SCTransform-normalized and the top 3000 highly variable genes across cells were selected. The datasets were integrated based on anchors identified between datasets before Principal Component Analysis (PCA) was performed for linear dimensional reduction. After normalization and scaling, a shared nearest neighbor (SNN) Graph was constructed to identify clusters on the low-dimensional space (top 30 statistically significant principal components, PCs). An unbiased clustering according to the recommendations of the Seurat package was used, if not provided by the authors. The cell types were assigned according to the authors’ recommendations or provided metadata. The alluvial plots were designed with the ggalluvial package (v0.12.5). For the DimPlots, the RNA-slot was used and every value above 0 was counted as “positive”. The differentially expressed markers were identified by the FindMarkers function (ident.1 was specified, and differences calculated to all other clusters) and the Wilcoxon signed-rank test. We used the SenMayo gene set (n=125) or secreted proteins, obtained by the human protein atlas, augmented by the SenMayo secreted SASP factors (n=1989^22^) to select the SASP factors or secreted proteins. The cytoscape bubble plot were designed with cytoscape (v3.1.0), and the size of each bubble is proportional to the avg_log2FC. The circular plots were designed with ggplot2 (v3.4.4). The spider plots were designed with the package fmsb (v0.7.5). The circle size is proportional to the log2FC compared to all other clusters, while the color codes the respective tissue. The bars show the median of all tissues in which the respective gene is upregulated.

For the intercellular communication heatmap, we first calculated the respective intercellular communication via CellChat (v1.6.1). The centrality values were extracted and used for the central heatmap for which we used GraphPad Prism (Version 9.0). The probability of the inferred communication was used for the sankey-network. The depiction of the sankey-network was done via the networkD3 package (v0.4) and exported via jsonlite (v1.8.7). Since muscle had too little p16+ cells for the CellChat analyses, these were excluded.

For the SCENIC analyses, we used the standard settings (v.1.3.1). For plotting the most important factors, the relative activity for each transcription factor above 1 was chosen for p21+ and p16+ cells, respectively, *per* tissue. Since liver has very few p16+ cells for proper calculation with SCENIC, these were excluded. For the pie charts, the sum of transcription factors sharing a tissue was calculated and the respective percentage is demonstrated within the plot.

### Cytometry by time of flight (CyTOF) analysis

The provided fcs-files were read into R by the flowCore package (v2.8.0) and transformed into a Seurat object. All subsequent analyses were following the standard Seurat procedure as described above.

### RNA-sequencing analysis

The fastq files were mapped to the murine genome (mm10), and analysis was performed using the DESeq2 package (v1.38.3) as previously described^13^. Significantly differentially regulated genes were selected by a Benjamini–Hochberg adjusted p value <0.05 and log2-fold changes above 0.5 or below −0.5.

### Statistics

Statistical analyses were performed using either GraphPad Prism (Version 9.0) or R version 4.2.0. A p-value <0.05 (two-tailed) was considered statistically significant.

### Code availability

The code for each figure will be provided by the first author upon reasonable request.

## Acknowledgements

This research was supported by the NIH through grants P01 AG062413 (to S.K., N.K.L., D.M, D.J., L.J.N., P.D.R. and J.F.P.), R01AG068048 (to J.F.P.), R01AG82708 (to J.F.P.), UG3CA268103 (to J.F.P.), R01 AG068182 (to D.J.), R01 AG063543 (L.J.N), Hevolution/AFAR (to D.J.), R01 AG076515 (to S.K, D.M., P.D.R., L.J.N.), U54 AG079754 (to S.K., N.K.L., P.D.R., L.J.N.), R01 AG055529 (to N.K.L), U54 AG076041 (to L.J.N., P.D.R.) and U19 AG056278 (to P.D.R., L.J.N.).

## Author contributions

All authors contributed to writing the manuscript and reviewed and approved of its submission for publication. D.S., S.K. and J.F.P conceived and directed the project. D.S. analyzed and interpreted the data with input from S.K. and J.F.P. M.L.D, N.K.L., P.D.R., L.J.N., D.J., R.L.K, D.G.M. contributed to the conceptual development of the project.

---

**Extended Data Figure 1.**
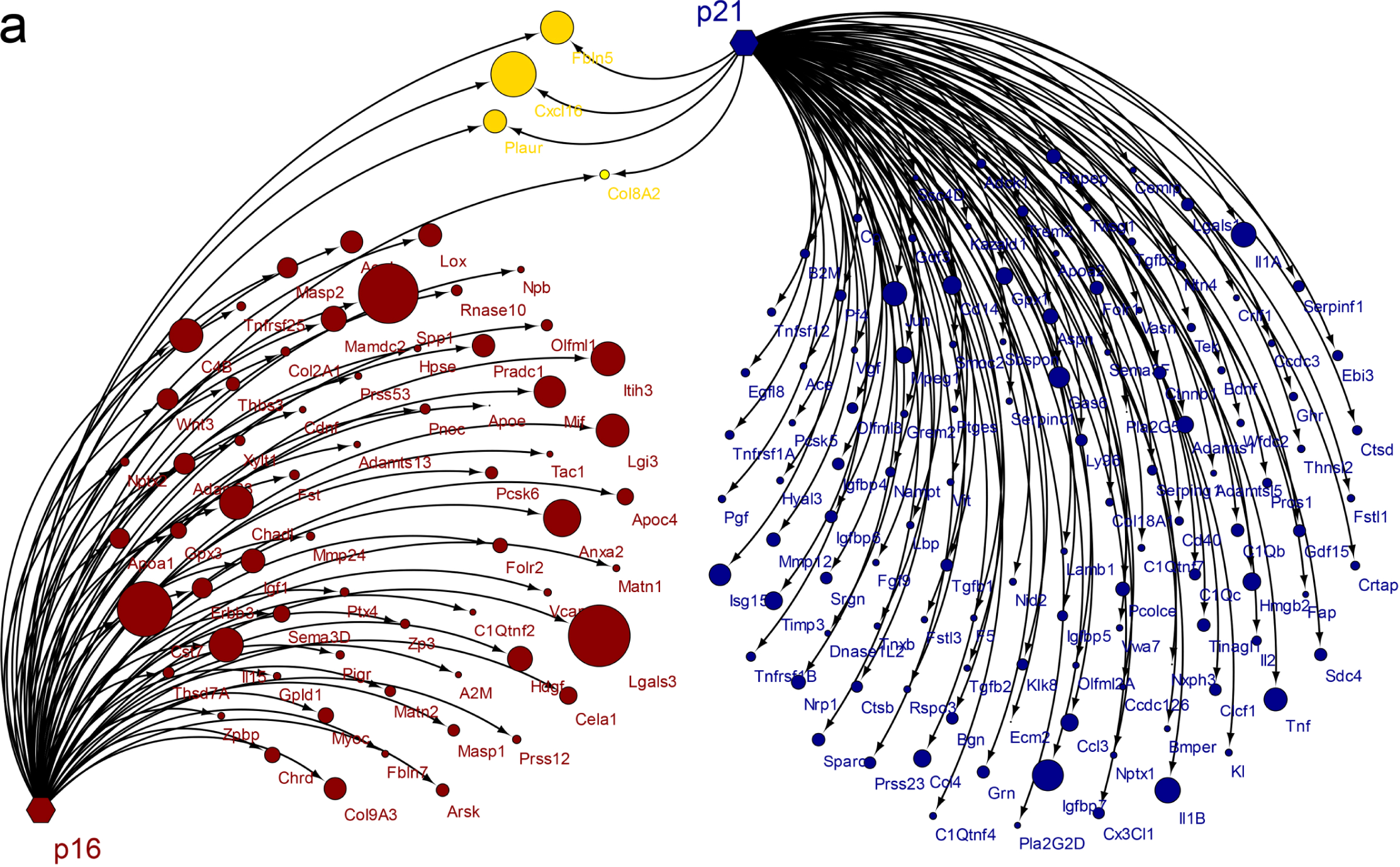
Comprehensive examination of the mRNA expression profiles for the whole secretome and SenMayo in aging murine hippocampus. (**a**) In the murine hippocampus, there is minimal overlap between the mRNA expression of the whole secretome + SenMayo in p21+ and p16+ cells.

**Extended Data Figure 2.**
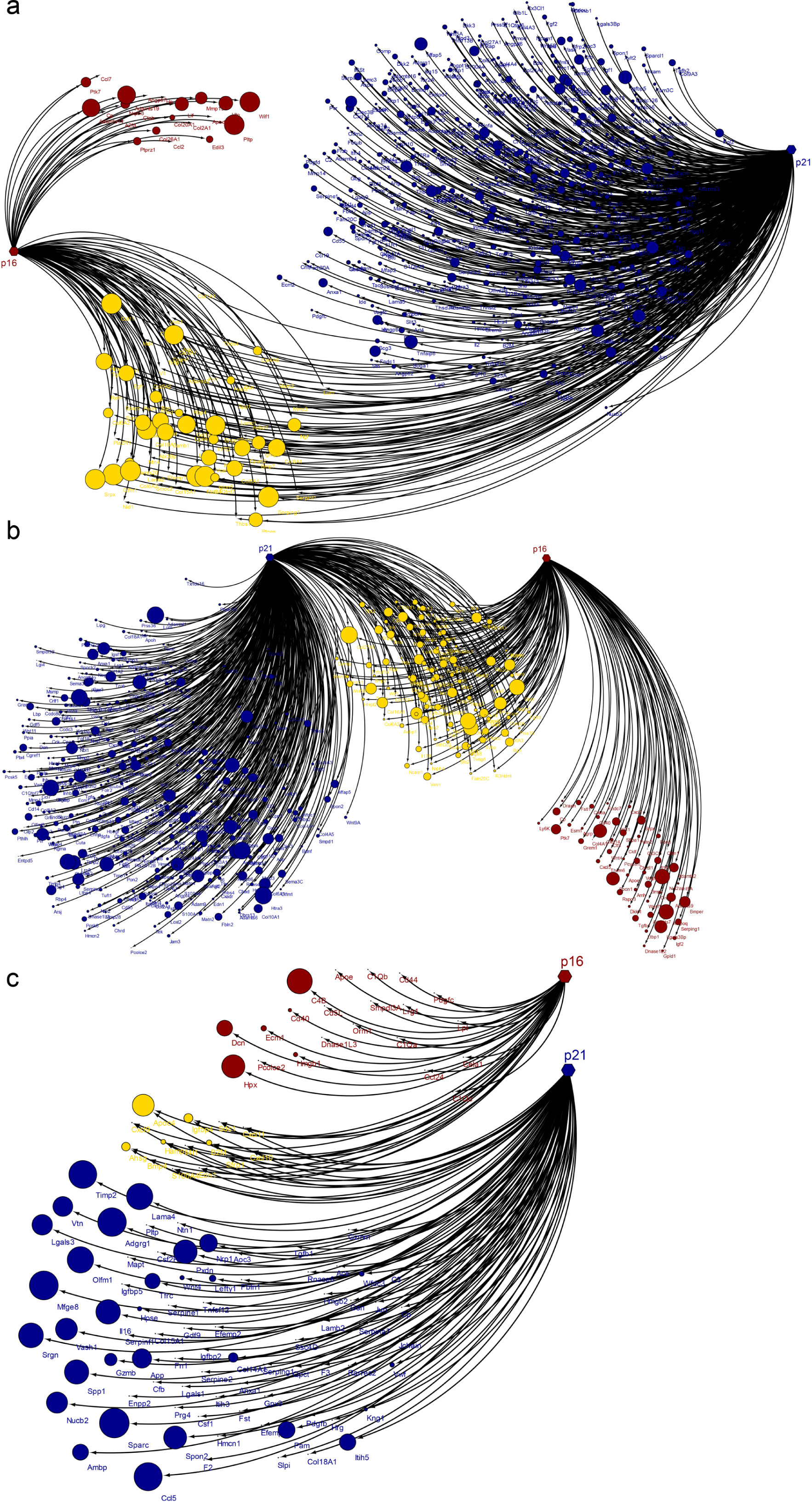
Comprehensive analysis of mRNA expression profiles for the whole secretome and SenMayo in aging murine muscle, bone, and liver. (**a**) In the muscle, there is some overlap with a noticeably larger p21-associated whole secretome and SenMayo mRNA expression compared to p16. (**b**) Similar to the muscle, in the bone, this phenomenon is comparable, with some overlap but a more pronounced p21-associated secretome and SenMayo. (**c**) Within the liver, there is minimal overlap between p21+ cells and p16+ cells, with the p21+ cells predominantly expressing the majority of factors.

**Extended Data Figure 3.**
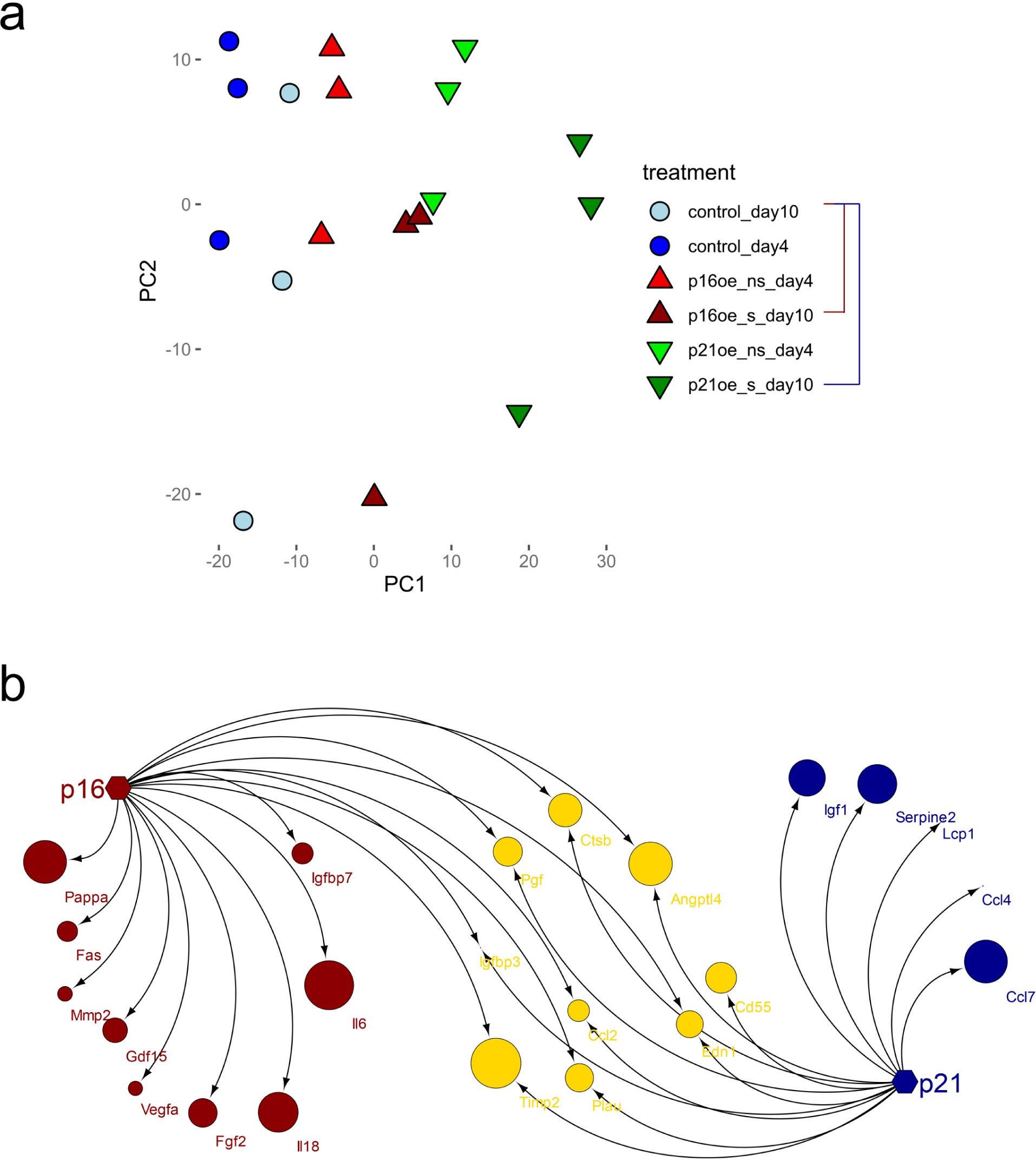
Overexpression of p16 and p21 reveals distinct SASP profiles (GSE117278). (**a**) A PCA plot illustrating control-MEFs on day 4 and 10, as well as p16-overexpressing cells (adeno-Cre-EGFP virus Ai14;L-p16 injection into the tail) on day 4 and 10, compared to p21-overexpressing cells (Ai14;L-p21) on day 4 and 10. (**b**) A comparable number of SASP factors are expressed in p16 (red)- vs. p21 (blue)-positive cells, with overlap indicated in yellow.

**Extended Data Figure 4.**
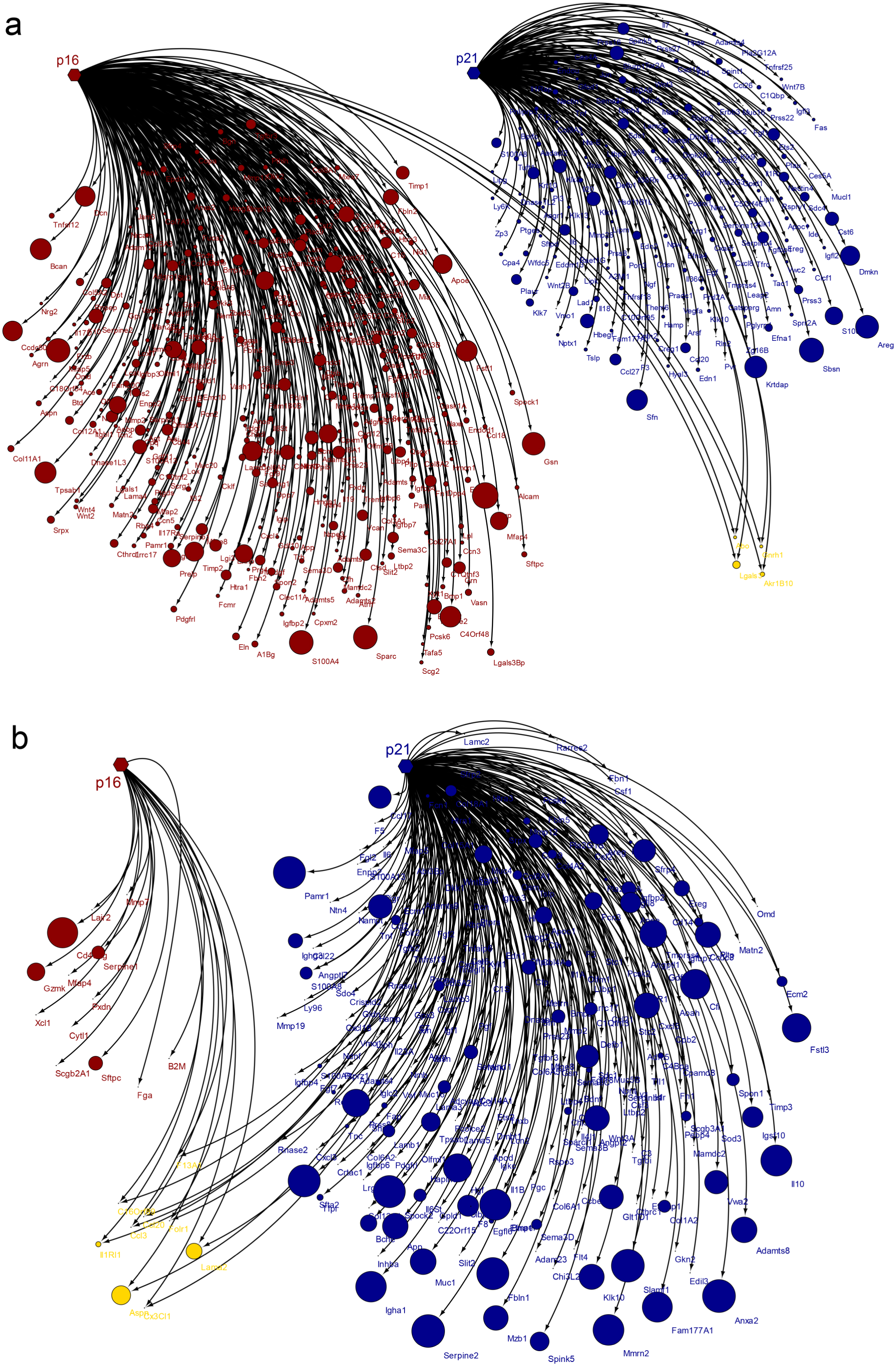
Analysis of the whole secretome and SenMayo in human skin and lung during aging. (**a**) In the skin, there is minimal overlap between p16- and p21-positive cells in terms of mRNA expression of the whole secretome and SenMayo. Unlike the majority of examined tissues, the mRNA expression of the entire secretome and SenMayo associated with p16 is as extensive as that associated with p21. (**b**) In the lung, the p21+ cells express a large number of secreted factors, with a very small overlap with p16+ cells.

**Extended Data Figure 5.**
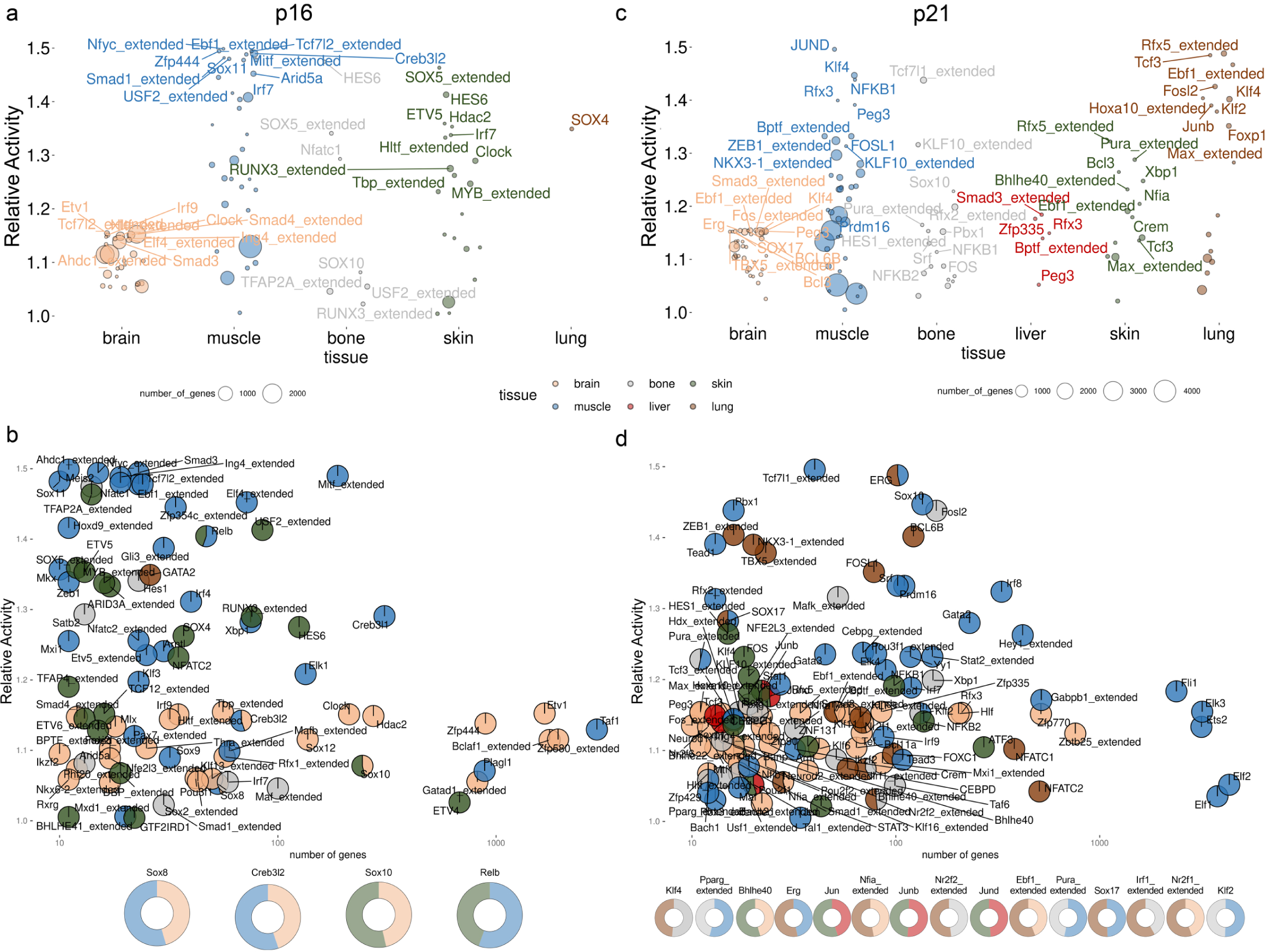
SCENIC analysis of the regulating transcription factors of p16+ and p21+ cells. (**a**) When analyzing five distinct tissues for regulatory factors in p16+ cells, each tissue exhibits a substantial array of transcription factors governing the behavior of p16+ cells. (Since liver has very few p16-positive cells for proper calculation with SCENIC, these were excluded). (**b**) The x-axis, represents the number of genes, plotted against the y-axis, illustrating the relative activity of the respective transcription factor, highlights the significance of each factor. Interestingly, only four factors (*Sox8*, *Creb3l2*, *Sox10*, and *Relb*) exhibit consistency across multiple tissues in the context of p16-positive cells. (**c**) In the p21+ cells, the regulating transcription factors show a high heterogeneity across tissues. (**d**) The transcription factors associated with p21 are predominantly specific to a single organ, but a few transcription factors such as *Erg*, *Sox17*, *Klf4*, *Jun*, *Klf2*, and others, regulate p21+ cells in two tissues.

